# Yin Yang 1-dependent TET2 regulation is essential for early T cell development

**DOI:** 10.64898/2026.01.27.701530

**Authors:** Yinghua Wang, Marjorie Brand, Matthew A. Nangle, Robert J. Lipinski, Xuan Pan

## Abstract

Yin Yang 1 (YY1) is a multifunctional transcription factor and mammalian Polycomb Group (PcG) protein critical for lymphocyte development. While YY1 is required for early T-cell development and survival, the underlying mechanisms remain incompletely defined. Herein, we utilize the YY1 REPO domain conditional knockout mouse model (*Yy1^-/ΔREPO^*) to further dissect the YY1-PcG domain dependent epigenetic regulation in early T cell development. *Yy1^-/ΔREPO^* mice show a developmental block at the double-negative (DN) 3 to DN4 T cell transition, with expansion of the DN3 population and reduced TCRβ^+^ DN4 T cells. The genetic network governing T cell differentiation is dysregulated in *Yy1^-/ΔREPO^* DN3 T cells. YY1 binds directly to the *Tet2* promoter, and deletion of the YY1 REPO domain leads to downregulation of the DNA demethylase TET2 in DN3 T cells. Although deletion of the YY1 REPO domain does not impair YY1 binding at the *Tet2* promoter, H3K4me3 enrichment at the promoter is reduced. Pharmacologic inhibition of TET catalytic activity in wild-type DN thymocytes partially recapitulates the developmental defects in *Yy1^-/ΔREPO^* DN thymocytes, whereas re-expression of TET2 catalytic domain in *Yy1^-/ΔREPO^* DN thymocytes partially rescues T cell development. Collectively, these findings reveal a previously unappreciated link between YY1 REPO domain-dependent regulation and TET2-mediated epigenetic control during early T-cell development.

## Introduction

The Ten-Eleven Translocation (TET) family proteins, TET1, TET2, and TET3, are Fe(II)- and α-ketoglutarate-dependent dioxygenases that catalyze the oxidation of 5-methylcytosine and promote DNA demethylation^1–4^. Through their regulation of DNA methylation and chromatin modification, TET proteins play critical roles in hematopoietic development, stem cell function, aging, and malignant transformation. Among the three family members, TET2 is particularly important in hematopoiesis, as loss-of-function mutations are frequently detected in clonal hematopoiesis of indeterminate potential (CHIP) and hematologic malignancies, including acute myeloid leukemia and T-cell lymphoma^5–10^.

Accumulating evidence has also established important roles for TET proteins in T cell biology. Combined deletion of TET2 and TET3 in double-positive (DP) thymocytes results in lymphoproliferative disease characterized by antigen-dependent expansion of invariant natural killer T (iNKT) cells^11^. In mature T cells, TET proteins regulate helper T-cell differentiation, maintain regulatory T-cell stability through demethylation of the FoxP3 locus, and shape CD8⁺ T-cell fate decisions during activation^12–16^. TET2 and TET3 have additionally been implicated in maintaining T-cell receptor (TCR) repertoire diversity during T-cell expansion^17,18^. Despite these advances, the roles of TET proteins during early thymocyte development, particularly at the double-negative (DN) stages, remain poorly understood.

Early T-cell development requires tightly coordinated epigenetic regulation, including dynamic changes in DNA methylation, DNA demethylation, and histone modifications that establish stage-specific gene expression programs^19–28^. Yin Yang 1 (YY1) is a multifunctional transcription factor, chromatin regulator, and Polycomb Group (PcG) protein that plays critical roles in lymphocyte development^29–38^. We previously demonstrated that YY1 is required for early T-cell development and Notch signaling, as hematopoietic-specific deletion of *Yy1* results in a developmental block at the DN3 stage^29,37^.

The PcG function of YY1 is mediated by its Polycomb Recruitment (REPO) domain (amino acids 201-226), which recruits PcG proteins, including the PRC2 component EZH2, to promote chromatin-mediated gene regulation^39–42^. Although deletion of the REPO domain does not impair YY1 DNA binding, transcriptional activation, or transient transcriptional repression, it abolishes YY1-dependent PcG functions, including Polycomb protein recruitment and stable epigenetic repression^41^. Using a retroviral bone marrow transplantation model, we previously found that YY1 regulates T-cell development through both REPO-dependent and REPO-independent mechanisms^37^. Specifically, YY1 control of DN1 transition and Notch signaling is independent of the REPO domain, whereas the REPO domain is required for early thymocyte survival^37^. However, the molecular mechanisms by which YY1 PcG function regulates early T-cell development remain unclear. To address this question, we utilized a YY1 REPO domain-deficient mouse model (*Yy1^f/ΔREPO^ Vav-Cre*) to define the role of YY1-mediated epigenetic regulation during early thymocyte development.

Herein, our study identifies YY1 as an upstream regulator of TET family member TET2 during DN3 T cell development. Deletion of the YY1 REPO domain/PcG function blocks DN3-to-SP T cell development and downregulates *Tet2* expression in DN3 T cells. Mechanistically, YY1 directly binds to the *Tet2* promoter, and deleting the YY1 REPO domain preserves YY1 binding but reduces H3K4me3 enrichment at this locus. *Yy1^-/ΔREPO^* DN3 T cells are highly proliferative with a deregulated transcriptional network governing T cell differentiation. *Yy1^-/ΔREPO^* mice have reduced TCRβ+ DN4 T cells and are more prone to apoptosis. Functionally, pharmacologic inhibition of TET catalytic activity in wild-type cells partially recapitulates some of the early T cell development defects observed in *Yy1^-/ΔREPO^* DN T cells. Furthermore, ectopic expression of a TET catalytic domain (CD) partially rescued the DP and CD8+ T cell differentiation in *Yy1^-/ΔREPO^* DN T cells. Our study revealed a distinct YY1 REPO-TET2 axis in early T cell development, highlighting a previously unrecognized regulatory mechanism with potential broader implications for immune development.

## Methods

### Mice

*Yy1^+/ΔREPO^* mice were generated by a germline deletion of one copy of the REPO domain by CRISPR/Cas9 gene editing^43^ and the *Yy1* promoter region and exon 1 are flanked by loxP sites in the second allele. *Yy1^f/f^* mice^44,45^ were crossed with *Vav-Cre* mice to generate heterozygous *Yy1^f/+^ Vav-Cre* mice. *Yy1^+/ΔREPO^*mice were crossed with *Yy1^f/f^* mice to generate *Yy1^f/ΔREPO^*mice. *Yy1^f/ΔREPO^* mice were then crossed with *Yy1^f/+^ Vav-Cre* mice to generate *Yy1^f/ΔREPO^ Vav-Cre* mice (**Figure 1A**). All mice were maintained in a C57BL/6 genetic background. Both male and female mice were used in this study, and animals were analyzed at 4–5 weeks of age unless otherwise specified. All experiments involving mice were approved by the Institutional Animal Care and Use Committee of the University of Wisconsin-Madison and conform to the appropriate regulatory standards.

**Figure 1.**
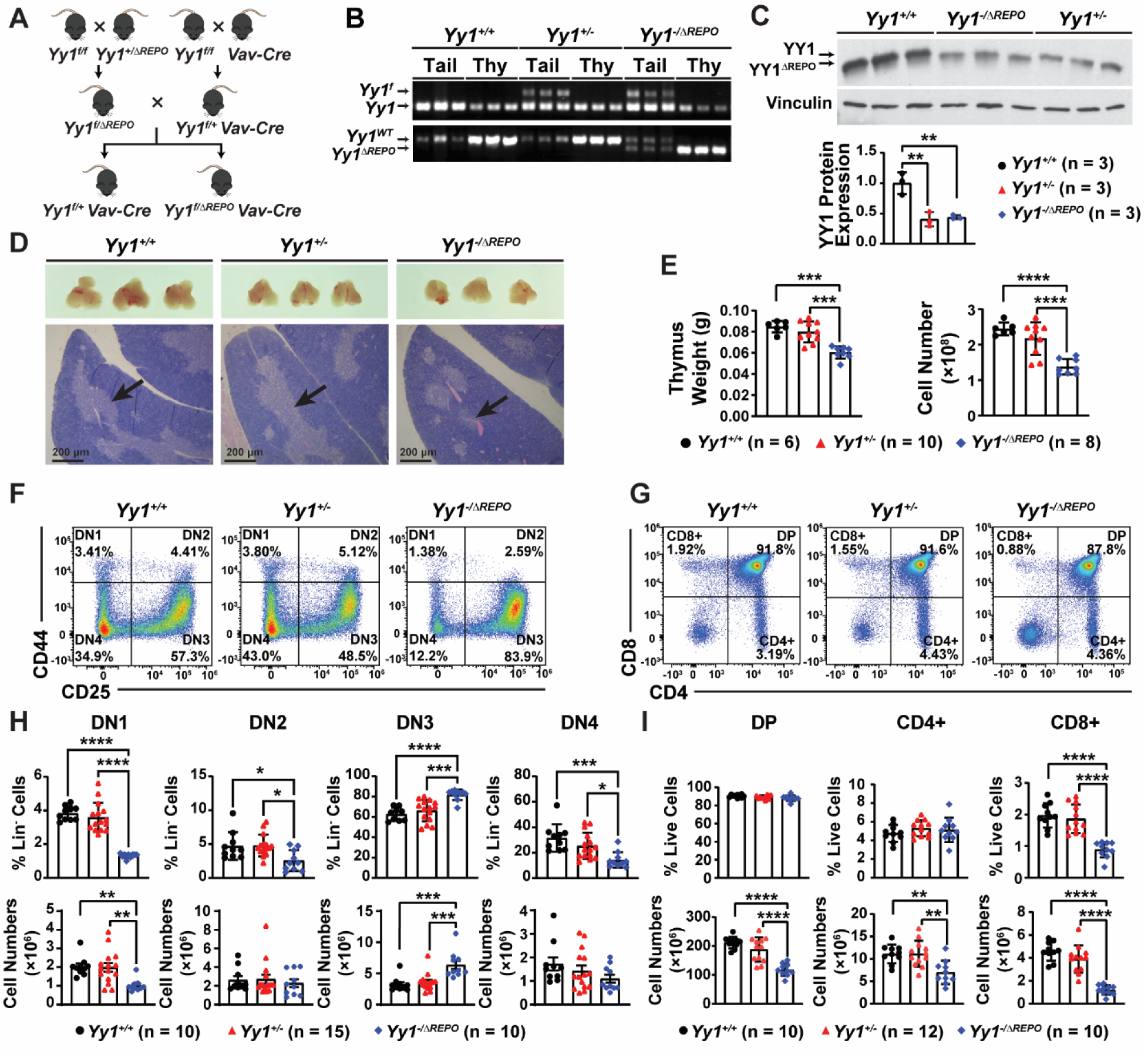
The YY1 REPO domain is critical for the DN3 to DN4 transition and for generating mature T cells in the mouse thymus. **(A)** Breeding strategy of *Yy1^f/ΔREPO^ Vav-Cre* mice. **(B)** PCR results of mouse tail samples and thymocytes from *Yy1^+/+^, Yy1^+/-^*, and *Yy1^-/ΔREPO^* mice. **(C)** Western blot showing wild-type YY1 and YY1^ΔREPO^ protein expression in total thymocytes. **(D)** Representative gross and histology images of the *Yy1^+/+^, Yy1^+/-^* and *Yy1^-/ΔREPO^*thymus. Arrows pointing to the medulla in the thymus. Histology pictures were taken with a 4X lens. **(E)** Quantification of weight and total cell numbers of thymus. **(F)** Representative flow gating strategy for DN1 (Lin^−^CD44^+^CD25^−^), DN2 (Lin^−^CD44^+^CD25^+^), DN3 (Lin^−^CD44^−^CD25^+^), and DN4 (Lin^−^CD44^−^CD25^−^) T cells. **(G)** Representative flow gating strategy for DP (CD4^+^CD8^+^), CD4^+^ T cells (CD4^+^CD8^−^), and CD8^+^ T cells (CD4^−^CD8^+^) T cells in thymus. (**H**) Quantification of percentage and absolute cell number of each DN population within Lin-thymocytes. **(I)** Quantification of percentage and absolute cell number of DP, CD4^+^, and CD8^+^ T cells. N represents the number of mice; data are presented as means ± SD; ∗*P* < .05, ∗∗*P* < .01, ∗∗∗*P* < .001, and ∗∗∗∗*P* < .0001 by one-way ANOVA.

### OP9-DL1 co-culture system

BM cells or thymocytes were isolated from *Vav-Cre*, *Yy1^f/+^ Vav-Cre*, and *Yy1^f/ΔREPO^ Vav-Cre* mice at 4 weeks of age. Lin^−^ BM cells were then isolated using the EasySep™ magnet system (Stemcell Technologies) with antibodies against CD4 (RM4-5), CD8 (53-6.7), B220 (RA3-6B2), CD11b (M1/70), NK1.1 (PK136), CD19 (eBio1D3), Ter 119 (TER-119), Gr1 (RB6-8C5), IgM (eB121-15F9), and CD127 (A7RS4) (eBioscience). Lin^−^ thymocytes were isolated by the same magnet system with antibodies against CD4 (RM4-5), CD8 (53-6.7), B220 (RA3-6B2, 1:400), CD11b (M1/70, 1:400), NK1.1 (PK136, 1:400), CD19 (eBio1D3, 1:400), Ter 119 (TER-119, 1:400) and Gr1 (RB6-8C5, 1:400) (eBioscience). Cells were plated on OP9-DL1^46^ feeder cells with MEM, FBS, penicillin/streptomycin, L-glutamine, Flt-3L (PeproTech), and IL7 (PeproTech). Cells were transferred to new OP9-DL1 feeder cells every 72 h. DN thymocytes were identified as lineage-negative cells lacking CD4, CD8, B220, TER119, Gr-1, Mac-1, and NK1.1 expression. DN T cells were further defined by CD44 (IM7) and CD25 (PC61.5) markers. For TET inhibition experiments, DMSO, BC339 (73 µM), or DMOG (0.5 mM) was added to the OP9-DL1 co-culture system for 72 hours.

### Retroviral infection

The FLAG-tagged TET2-CD^47,48^ was cloned into the GFP-expressing MigR1 vector. Lin^-^thymocytes were cultured overnight in OP9-DL1 coated 12-well plates, and spin infected with MigR1-TET2-CD or MigR1 retroviral supernatant containing polybrene (4 µg/ml). Transduction efficiency was assessed by GFP expression using flow cytometry 48 hours after the first infection.

### Compiling YY1 ChIP-seq data

YY1 binding in human CD4+ T cells (GSE25674; GRCh38/hg38) and mouse HSPC7 (GSE261032; GRCm38/mm10) were analyzed by using Integrative Genomics Viewer. GRCh38/hg38 was used for human datasets and mm10/GRCm38 was used for mouse datasets. YY1 binding at the *Tet* promoter regions was defined by overlap between processed BED peak intervals and the annotated transcription start site region.

### CUT&Tag qPCR

The Cleavage Under Targets And Tagmentation (CUT&Tag) was performed using Lin^-^ thymocytes and antibodies against YY1 (61779; Active Motif), H3K4me3 (ab213224; Abcam), and IgG (10500C; Invitrogen) as previously described^36,43^. Briefly, Lin^−^ thymocytes were bound to concanavalin A–coated magnetic beads, permeabilized, and incubated with primary antibodies. After incubation with secondary antibody, samples were treated with pA-Tn5 transposase to tagment antibody-bound chromatin regions. Tagmented DNA was purified and amplified to generate CUT&Tag libraries. CUT&Tag libraries were purified using QIAquick Gel Extraction Kit (Qiagen). Enrichment at target gene promoter regions was assessed by qPCR. IgG served as the background control, and fold enrichment was calculated as 2^(Ct_IgG_ – Ct_Target_).

## Statistical analysis

All data are expressed as mean ± SD. All statistical analyses were conducted using GraphPad Prism 10. Differences between the groups were determined using one-way ANOVA followed by Tukey’s post hoc test when comparing 3 or more groups. Two-way ANOVA followed by Tukey’s post hoc test was used for apoptosis assay and cell cycle analysis. P-values < 0.05 were considered statistically significant.

## Results

### The YY1 REPO domain is essential for DN3 to DN4 transition during early T cell development

To study YY1-dependent PcG regulation of early T cell development, *Yy1^f/ΔREPO^* mice were generated by CRISPR/Cas9 gene editing as previously described^43^. *Yy1^f/ΔREPO^*mice were crossed to the *Vav-Cre* system (**Fig. 1A**). In *Yy1^f/ΔREPO^ Vav-Cre* mice, one allele of YY1 was deleted upon the Cre recombinase expression specifically in the hematopoietic system at E11.5 of embryonic development^49^. The YY1 REPO domain was germline deleted at the second allele. PCR using thymus and tail genomic DNA confirmed hematopoietic-specific deletion of YY1 and germline deletion of the YY1 REPO domain in *Yy1^f/ΔREPO^ Vav-Cre* (*Yy1^-/ΔREPO^*) thymic cells (**Fig. 1B**). Western blot further supports that one allele of YY1 was deleted, leaving another allele of *Yy1^-/ΔREPO^* (**Fig. 1C**). In our study, it is essential to include *Yy1^f/+^ Vav-Cre* (*Yy1^+/-^*) as an additional control to rule out the impact of YY1 heterozygosity.

The *Yy1^-/ΔREPO^* mice have smaller thymus size, lower thymus weight, and fewer absolute thymocyte numbers compared with *Vav-Cre* (*Yy1^+/+^*) and *Yy1^+/-^* mice (**Figs. 1D and 1E**). Compared with *Yy1^+/+^* and *Yy1^+/-^* thymus, the *Yy1^-/ΔREPO^* thymus showed reduced medulla, a primary site for SP T cells maturation (**Fig. 1D**)^50,51^. Consistently, flow cytometry analysis revealed that *Yy1^-/ΔREPO^* thymus has reduced numbers of DP, CD4+, and CD8+ T cells (**Fig. 1G and 1I**). Within DN compartments, *Yy1^-/ΔREPO^* mice show decreased DN1, DN2, and DN4 percentages (**Figs. 1F and 1H**). In contrast, there was an increase of both percentage and absolute number of DN3 T cells in *Yy1^-/ΔREPO^* compared with *Yy1^+/+^* and *Yy1^+/-^* mice (**Figs. 1F and 1H**). Together, our results demonstrate that deletion of the YY1 REPO domain leads to a developmental block at the DN3 to DN4 stage in early T development and thymic atrophy. Because this phenotype was not observed in YY1 heterozygous mice, these data suggest that the YY1-mediated DN3 to DN4 transition requires its REPO / PcG domain.

### Genetic networks governing T cell differentiation are deregulated in the YY1 REPO domain deleted DN3 T cells

To further dissect the underlying mechanisms, bulk RNA-seq was performed in *Yy1^+/+^*, *Yy1^+/-^*, and *Yy1^-/ΔREPO^* DN3 (Lin^-^CD25^+^CD44^-^) T cells. *Yy1^-/ΔREPO^* has 1118 genes downregulated and 630 genes upregulated compared with *Yy1^+/+^* DN3 T cells, and 1231 genes downregulated and 328 genes upregulated compared with *Yy1^+/-^* DN3 T cells (**Fig. 2A**). To identify key molecular pathways altered in *Yy1^-/ΔREPO^* DN3 T cells, we performed Gene Set Enrichment Analysis (GSEA). Among the significantly deregulated GO M5 pathways, pathways associated with early T cell development, T cell activation, and T lineage commitment were downregulated in *Yy1^-/ΔREPO^* DN3 T cells compared to *Yy1^+/+^* (**Fig. 2B**). Furthermore, T cell differentiation and αβ T cell differentiation pathways, positive T cell selection, and T cell differentiation are deregulated in *Yy1^-/ΔREPO^* DN3 T cells (**Fig. 2C**). Notably, several downregulated genes in *Yy1^-/ΔREPO^* DN3 T cells have established roles in T cell signaling, survival, or differentiation (**Fig. 2D**). *Gimap3* has been implicated in T cell survival^52,53^; *Txk* contributes to TCR signaling and thymocyte selection^54,55^; and *Slamf6* participates in SLAM-family receptor signaling during NKT lineage development and T cell exhaustion^56,57^ (**Fig. 2D**). Several transcriptional regulators were reduced in *Yy1^-/ΔREPO^* DN3 T cells, including *Zbtb16*, which controls NKT cell development^58–60^; *Ep300,* which encodes the transcriptional coactivator p300 required for DP T cell development^61^; and *Ncor1*, which supports thymocyte survival and selection^62,63^ (**Fig. 2D**). Together, these transcriptomic changes indicate that loss of the YY1 REPO domain disrupts the gene regulatory network required for the TCR signaling, T cell survival and differentiation.

**Figure 2.**
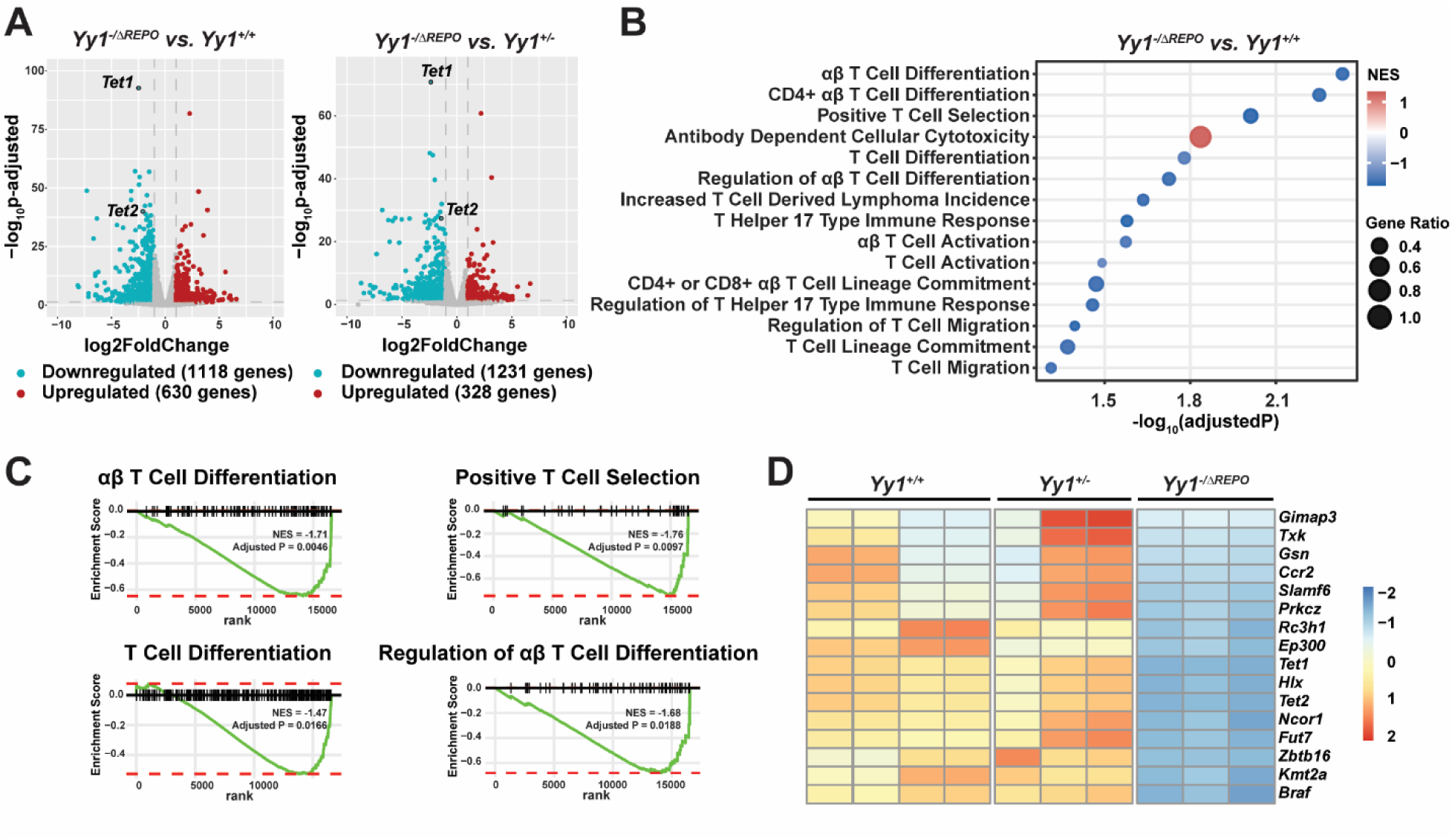
Deleting the YY1 REPO domain leads to dysregulation of T cell differentiation pathways in DN3 T cells. **(A)** Volcano plot of DEGs in *Yy1^-/ΔREPO^* DN3 T cells compared with *Yy1^+/+^* and *Yy1^+/-^* controls. **(B)** Enriched GSEA GO M5 pathways in *Yy1^-/ΔREPO^* DN3 T cells compared with *Yy1^+/+^* controls. Dot size represents the ratio of leading-edge genes contributing to the enrichment signal for each pathway. Color gradient indicates the normalized enrichment score (NES). **(C)** Deregulated T cell development pathways in *Yy1^-/ΔREPO^* DN3 T cells compared to *Yy1^+/+^* control. **(D)** Heatmap showing downregulated genes associated with T cell differentiation in *Yy1^-/ΔREPO^* DN3 T cells compared to *Yy1^+/+^* and *Yy1^+/-^* controls.

### The YY1 REPO domain / PcG function is essential for TET2 expression in DN3 T cells

Among the top differentially expressed genes, the DNA demethylase TET family members *Tet1* and *Tet2* mRNA level were downregulated in *Yy1^-/ΔREPO^* DN3 T cells compared with *Yy1^+/+^* and *Yy1^+/-^* cells (**Figs. 2A and 3A**). Although *Tet1* was differentially expressed, its absolute expression in DN3 thymocytes was low compared with *Tet2* and *Tet3*, consistent with previous reports about relatively low expression of *Tet1* in mouse thymocytes and peripheral T cells^11,64^. Western blot analysis further confirmed a reduced TET2 protein expression in *Yy1^-/ΔREPO^* DN T cells compared with *Yy1^+/+^* and *Yy1^+/-^* samples (**Fig. 3B**). Our study supports that the YY1 REPO domain deletion leads to a reduced TET2 expression in mouse DN T cells.

**Figure 3.**
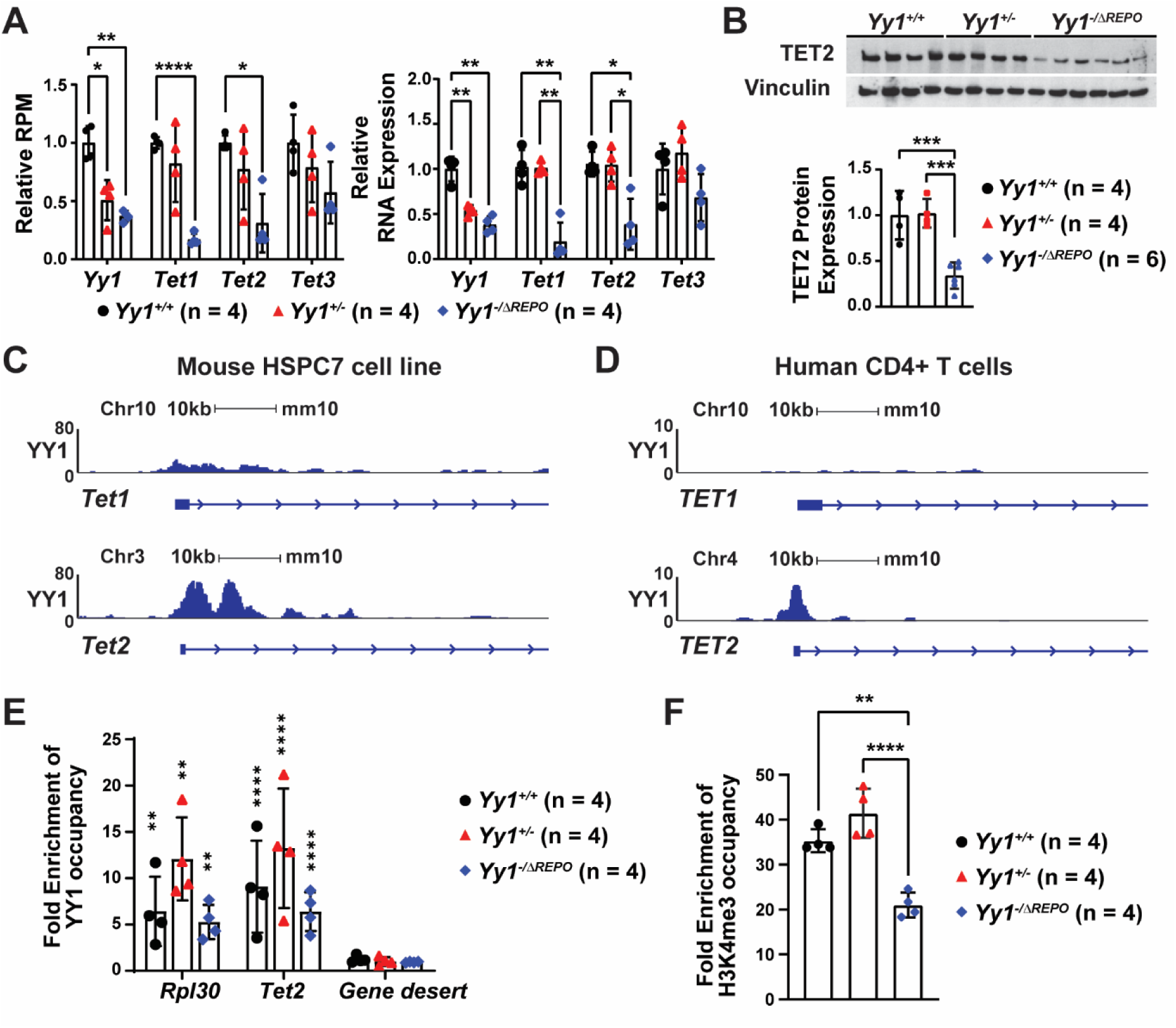
The YY1 REPO domain regulates *Tet1* and *Tet2* expressions in DN3 T cells. **(A)** *Yy1, Tet1, Tet2,* and *Tet3* transcript levels in RNA-seq and qRT-PCR validations. **(B)** Western blot shows TET2 protein expression level and quantification in DN T cells. **(C-D)** YY1 binding peaks at *TET1* and *TET2* promoter areas in mouse HSPC-like cell line HSPC7 **(C)** and human CD4^+^ T cells **(D)**. **(E)** CUT&Tag qPCR showing YY1 binding at the *Tet2* promoter in Lin^-^ thymocytes, compared to the gene desert negative control. *Rpl30* serves as a positive control for YY1 binding. **(F)** CUT&Tag qPCR showing enrichment of H3K4me3 at the *Tet2* promoter area in Lin^-^thymocytes. N represents the number of mice in **3A**, **3B**, and **3E**; N represents technical replicates in **3F**; data are presented as means ± SD; ∗∗*P* < .01, ∗∗∗*P* < .001, and ∗∗∗∗*P* < .0001 by two-way ANOVA (**A** and **E**) and one-way ANOVA (**B** and **F**).

To assess whether YY1 can directly regulate *Tet2* gene expression, we first evaluated YY1 binding at the *Tet2* promoter by utilizing our previously published CUT&Tag data set in mouse hematopoietic stem and progenitor cells (HSPC)-like cell line HSPC7 (GSE261032). Our data supports YY1 occupancy at the *Tet2* promoter in HSPC7 cells (**Fig. 3C**). Next, we compiled previously published ChIP-seq datasets and showed YY1 binding at the *TET2* promoter, but not at the *TET1* promoter in human CD4^+^ T cells (GSE25674) (**Fig. 3D**). Furthermore, by qPCR amplification of DNA from CUT&Tag library generated from Lin-thymocytes from *Yy1^+/+^*, *Yy1^+/-^*and *Yy1^-/ΔREPO^* mice, we confirmed YY1 binding at the *Tet2* promoter in those samples (**Fig. 3E**). Interestingly, although deletion of the YY1 REPO domain reduced TET2 expression (**Figs. 3A and 3B**), YY1 occupancy at the *Tet2* promoter remained unchanged (**Fig. 3E**). Notably, *Yy1^-/ΔREPO^* DN3 T cells exhibited reduced H3K4me3 histone enrichment at the *Tet2* promoter compared to *Yy1^+/+^* and *Yy1^+/-^* controls (**Fig. 3F**). Together, our data revealed a distinct YY1-TET2 axis in early T cell development. The YY1 REPO domain is essential for TET2 expression and is required to maintain the active chromatin mark at the *Tet2* promoter in DN T cells.

To further evaluate the effects of the YY1 REPO domain deletion on global DNA methylation status, we assessed the levels of 5-methylcytosine (5-mC) and 5-hydroxymethylcytosine (5-hmC) in *Yy1^+/+^*, *Yy1^+/-^*, and *Yy1^-/ΔREPO^* DN3 T cells. There were no statistically significant differences detected in global 5-mC or 5-hmC levels in *Yy1^+/+^*, *Yy1^+/-^*, and *Yy1^-/ΔREPO^* DN3 T cells **(Figs. S1A and S1B)**. Dot blot also confirmed no statistically significant differences in global 5-hmC levels in *Yy1^-/ΔREPO^* Lin^-^ thymocytes (**Figs. S1C and S1D**). These findings suggest that while TET2 is downregulated, global DNA 5-mC and 5-hmC remain unchanged in *Yy1^-/ΔREPO^* DN3 T cells, consistent with prior reports that loss of TET2 does not typically cause global hypermethylation^65,66^. TET2-mediated demethylation may occur in a locus-specific manner, and there may be functional redundancy among TET family members, particularly retained TET3 activity ^12,67,68^.

### The YY1 REPO domain is essential for TCRβ expression and survival during early T cell development

During DN3 T cell development, only cells that successfully rearrange the TCRβ and express pre-TCR can pass the developmental checkpoint and progress to DN4 T cells^69,70^. We examined the surface expression of TCRβ, a component of the pre-TCR complex by flow cytometry analysis ^71^. *Yy1^-/ΔREPO^* DN4 T cells showed a modest but significant reduction of TCRβ^+^ cell percentage and absolute numbers in contrast to no changes in DN3 T cells (**Figs. 4A and 4B**).

**Figure 4.**
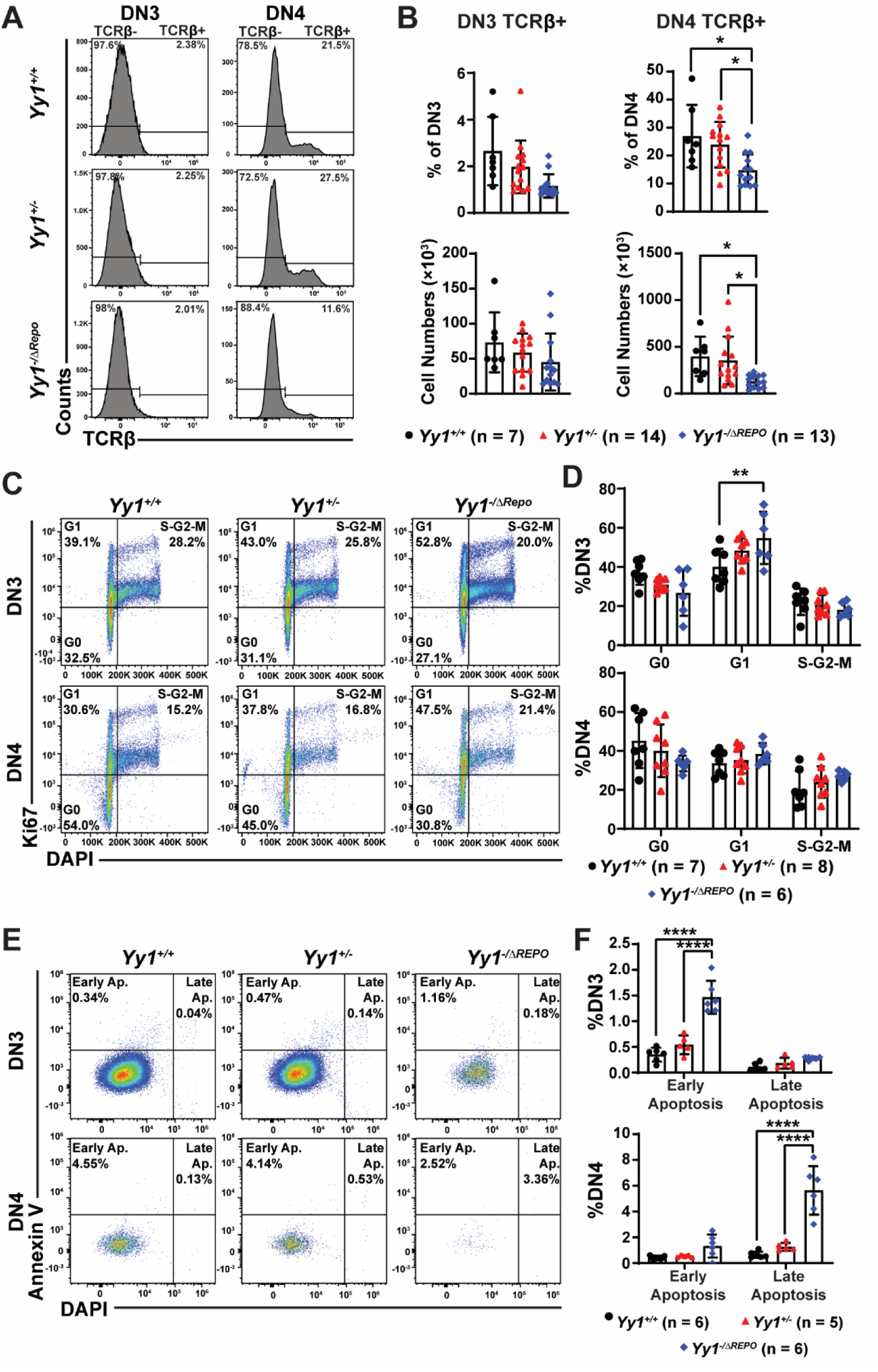
Deleting the YY1 REPO domain leads to impaired TCRβ cell surface expression, hyperproliferation, and reduced survival in DN T cells. **(A)** Representative flow plots showing TCRβ expression in *Yy1^+/+^*, *Yy1^+/-^*, and *Yy1^-/ΔREPO^* DN3 and DN4 T cells. **(B)** Quantification of percentages and cell numbers of TCRβ^+^ cells in DN3 and DN4 T cells. **(C)** Representative flow plots of the Ki67/DAPI proliferation assay in DN3 and DN4 T cells. **(D)** Quantification of G0 (Ki67^-^DAPI^-^), G1 (Ki67^+^DAPI^-^) and S/G2/M (Ki67^+^DAPI^+^) cells in DN3 and DN4 T cells. **(E)** Representative flow plots of apoptosis assays in OP9-DL1 co-cultured DN3 and DN4 cells. **(F)** Quantification of early apoptotic (Annexin V^+^, DAPI^-^) and late apoptotic (Annexin V^+^, DAPI^+^) cells in OP9-DL1 co-cultured DN3 and DN4 cells. N represents the number of mice; data are presented as means ± SD; ∗*P* < .05, ∗∗*P* < .01, and ∗∗∗∗*P* < .0001 by one-way ANOVA (**B**) and two-way ANOVA (**D** and **F**).

As DN3 T cells with successfully rearranged TCRβ undergo clonal expansion and receive survival signals to further differentiate into DN4 T cells^72–74^, we next assessed the cell proliferation and survival. The Ki67 proliferation assay revealed that *Yy1^-/ΔREPO^* DN3 T cells have a higher percentage of cells in the G1 phase (**Figs. 4C and 4D**), supporting a hyperproliferative state in these cells. Although *Yy1^-/ΔREPO^* DN3 and DN4 T cells did not show survival defects *in vivo* (**Figs. S2A-B**), apoptosis is challenging to detect in vivo due to the active phagocytosis mediated by macrophages in the thymus^75,76^. Thus, we tested apoptosis *in vitro* using the OP9-DL1 differentiation system. In this system, lineage-negative bone marrow cells were co-cultured with OP9 cells expressing the Notch ligand DL1, which supports T lineage differentiation in the presence of Flt3-L and IL-7 cytokines (**Fig. S2C**)^77^. In the OP9 co-culture system, *Yy1^-/ΔREPO^* DN3 and DN4 T cells were more apoptotic and failed to survive (**Figs. 4E and 4F**). Thus, deleting the YY1 REPO domain is essential for proper TCRβ expression and cell survival during early T cell development and deletion of the YY1 REPO domain leads to hyperproliferative DN3 T cells.

### Pharmacological inhibition of TET catalytic activity leads to early T cell developmental defects

To evaluate whether YY1 regulation of TET family proteins is a key driver for DN T cell development, we pharmacologically inhibited TET catalytic functions by using Bobcat339 (BC339) or Dimethyloxalylglycine (DMOG) in the OP9-DL1 co-culture system (**Fig. 5A**). BC339, a cytosine derivative, inhibits TET2 enzymatic activity by binding to its catalytic pocket and blocking the oxidation of 5-mC^78^. DMOG is a 2-oxoglutarate analog that broadly inhibits α-ketoglutarate–dependent dioxygenases, including TET enzymes^79,80^. In OP9-DL1 co-culture system, *Yy1^-/ΔREPO^* Lin^-^ thymocytes showed increased percentages of DN3 T cell (**Fig. S3A**), and reduced percentages and numbers of DP and SP T cell compared with *Yy1^+/+^* and *Yy1^+/-^*samples (**Fig. S3B**). Thus, deletion of the YY1 REPO domain in OP9-DL1 co-culture system partially recapitulates the DN developmental block detected in *Yy1^-/ΔREPO^* mice (**Figs. 1F and 1I**), supporting its use for further functional analyses. At 72h after BC339 treatment, *Yy1^+/+^* and *Yy1^+/-^*Lin^-^ thymocytes show reduced total cell numbers (**Fig. 5B**) and increased percentages of DN3 T cells (**Fig. 5C**). BC339 treatment did not significantly alter proliferation (**Fig. 5D**). However, BC339 treatment increased percentages of early apoptotic cells in *Yy1^+/+^*, *Yy1^+/-^* and *Yy1^-/ΔREPO^* DN3 T cells compared with corresponding DMSO-treated controls (**Fig. 5E**), consistent with the survival defects observed in *Yy1^-/ΔREPO^* DN3 T cells (**Fig. 4F**). Following reduced DN3 T cell numbers (**Fig. 5C**), BC339 treatment reduced the percentage and absolute number of DP, CD4+, and CD8+ T cells compared to DMSO-treated control cohorts (**Fig. 5F**). Thus, inhibiting TET catalytic activity in wild-type thymocytes partially recapitulates DN T development defects detected in *Yy1^-/ΔREPO^* mice. As *Yy1^-/ΔREPO^* DN3 thymocytes already had reduced TET2 expression (**Fig. 3B**), further pharmacologic inhibition of TET catalytic activity produced limited developmental effects in *Yy1^-/ΔREPO^* cohort. There were no significant changes in DN3 T cells percentages and no change in DP and SP T cell number or percentage in *Yy1^-/ΔREPO^* Lin^-^ thymocytes treated with BC339 (**Figs. 5C and 5F**). Similar results were also observed in cells treated with DMOG (**Fig. S4**). Together, these data support that inhibition of TET catalytic functions in wild-type Lin^-^ thymocytes impairs the DN3 to DN4 T cell transition and promotes DN3 apoptosis, thereby recapitulating key defects observed in the *Yy1^-/ΔREPO^* DN T cells.

**Figure 5.**
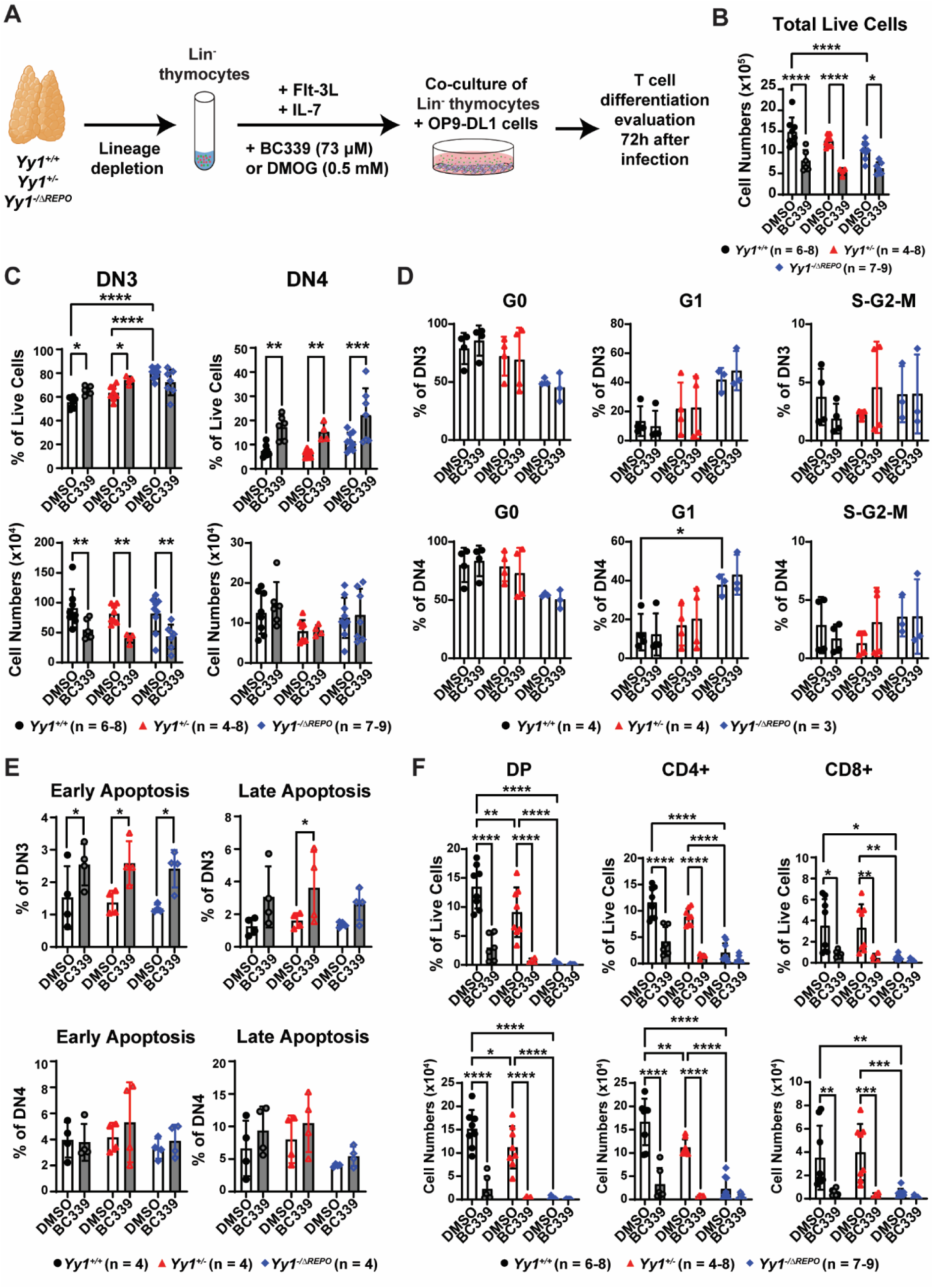
Pharmacological inhibition of TET catalytic activity leads to early T cell developmental defects observed in *Yy1^-/ΔREPO^* T cells. **(A)** Schematic diagram of OP9-DL1 co-culture with *Yy1^+/+^, Yy1^+/-^*, and *Yy1^-/ΔREPO^*Lin^-^ thymocytes treated with TET inhibitors. **(B)** Total cell counts of Lin^-^ *Yy1^+/+^, Yy1^+/-^*, and *Yy1^-/ΔREPO^* thymocytes co-cultured with OP9-DL1 treated with TET inhibitor BC339 or DMSO control. **(C)** Quantification of co-cultured DN3 and DN4 T cells treated with TET inhibitor BC339 or DMSO control. **(D)** Quantification of G0 (Ki67^-^DAPI^-^), G1 (Ki67^+^DAPI^-^) and S/G2/M (Ki67^+^DAPI^+^) cells in DN3 and DN4 T cells treated with BC339 or DMSO control in OP9-DL1 co-culture system. **(E)** Quantification of early apoptotic (Annexin V^+^, DAPI^-^) and late apoptotic (Annexin V^+^, DAPI^+^) cells in DN3 and DN4 T cells treated with BC339 or DMSO control in OP9-DL1 co-culture system. **(F)** Quantification of co-cultured DP, CD4+, and CD8+ T cells treated with TET inhibitor BC339 or DMSO control. N represents the number of mice; data are presented as means ± SD; ∗*P* < .05, ∗∗*P* < .01, ∗∗∗*P* < .001, and ∗∗∗∗*P* < .0001 by two-way ANOVA.

### Expression of a TET2 CD partially rescued T cell development ex vivo

Next, we tested whether restoring TET2 catalytic activity in the *Yy1^-/ΔREPO^* Lin^-^ thymocytes can rescue DN3 T cell development. Lin^-^ thymocytes were retrovirally transduced with MigR1-TET2-CD^47,48^ or MigR1 empty vector control and co-cultured with OP9-DL1 feeder cells^37^ (**Fig. 6A**). Compared with MigR1 empty vector controls, MigR1-TET2-CD transduction significantly increased *TET2-CD* mRNA expression in *Yy1^-/ΔREPO^* Lin^-^ thymocytes (**Fig. S5A**). In the OP9-DL1 co-culture system, *Yy1^-/ΔREPO^* Lin^-^ thymocytes transduced with empty vector showed impaired T cell differentiation with higher percentages of DN3 T cells and fewer DP, CD4+, and CD8+ T cells compared with *Yy1^+/+^* and *Yy1^+/-^* controls (**Figs. 6B and S5B**), consistent with the developmental block observed in *Yy1^-/ΔREPO^* mice (**Figs. 1F-I**). Ectopic expression of TET2-CD in *Yy1^-/ΔREPO^* cells showed decreased percentages of DN3 T cells and increased percentages of DP and CD8+ T cells compared with empty vector–transduced *Yy1^-/ΔREPO^* cells (**Fig. 6B**). These results support that restoration of TET2 catalytic activity partially rescues the early T cell development block caused by the YY1 REPO deletion. Together, these findings highlight a critical role of YY1–TET2 axis in T cell development.

**Figure 6.**
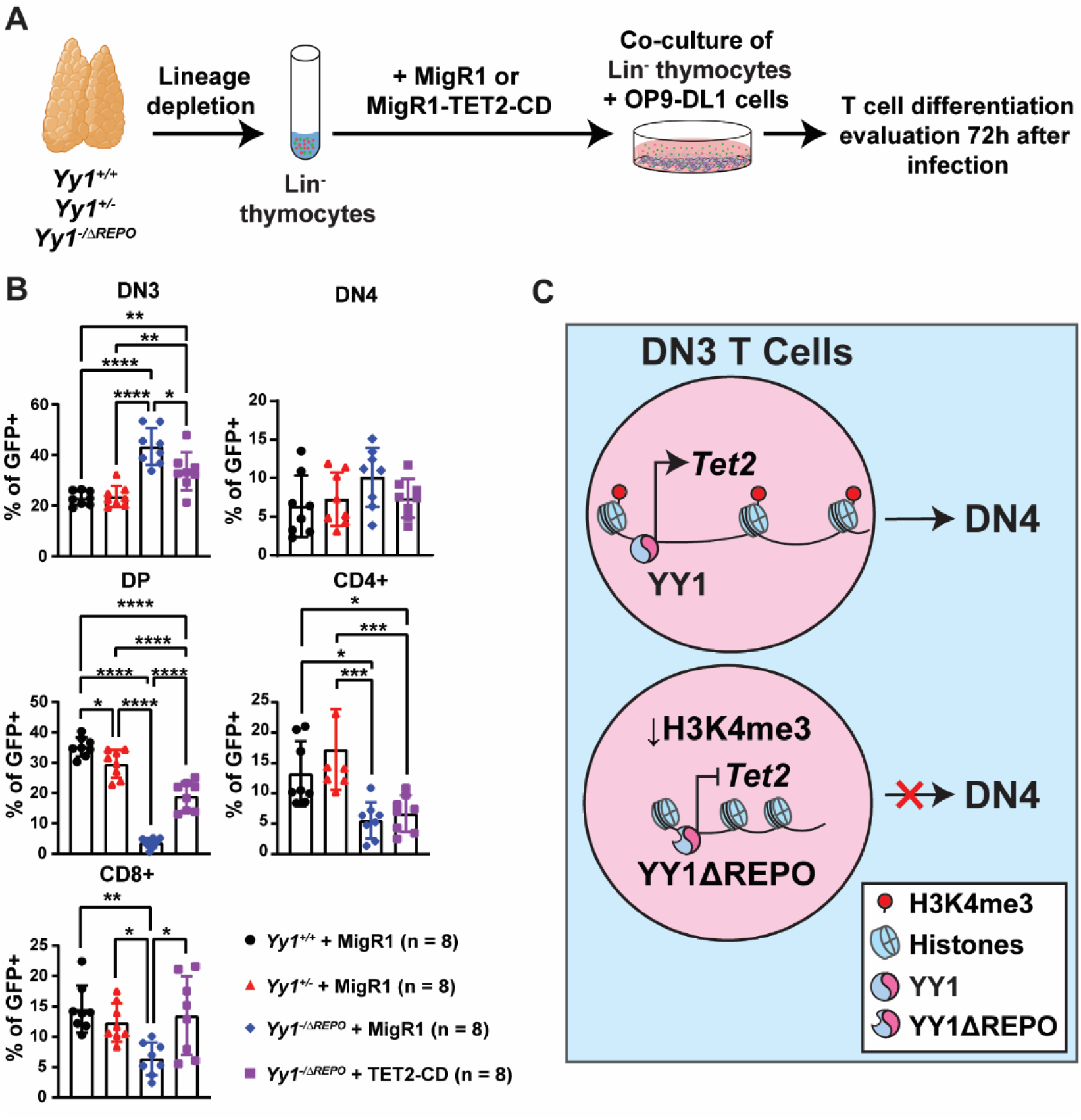
Ectopic expression of a TET2 catalytic domain partially rescues early T cell development. **(A)** Schematic diagram of Lin^-^ thymocytes retrovirally transduced with MigR1-TET2-CD or MigR1 vector control and cultured with OP9-DL1 for T cell differentiation. **(B)** Quantification of percentages of DN3, DN4, DP, CD4+ and CD8+ T cells. Cells infected with MigR1 or MigR1-TET2-CD are gated on GFP+. **(C)** A schematic model: when the full-length YY1 binds at the *Tet2* promoter, it promotes TET2 expression in DN3 T cells and supports DN3 to DN4 T cell transition. Deletion of the YY1 REPO domain leads to reduced H3K4me3 enrichment at the *Tet2* promoter, a less transcriptionally permissive chromatin state, and reduced *Tet2* expression. N represents the number of mice. Data are presented as means ± SD; ∗*P* < .05, ∗∗*P* < .01, ∗∗∗*P* < .001, and ∗∗∗∗*P* < .0001 by one-way ANOVA.

## Discussion

TET2 is a key epigenetic regulator of hematopoiesis and hematologic malignancy, yet its role during early T-cell development remains incompletely defined^5–10^. In this study, we identify a previously unrecognized YY1 REPO domain–TET2 regulatory axis that contributes to DN3 thymocyte differentiation. Using a CRISPR/Cas9-generated *Yy1^−/ΔREPO^*mouse model, we demonstrate that loss of the YY1 REPO domain results in a developmental block at the DN3 stage (**Figs. 1F and 1H**), impaired generation of TCRβ+ DN4 cells (**Fig. 4B**), and reduced DP and SP thymocyte output (**Figs. 1G and 1I**). These defects are accompanied by dysregulation of the transcriptional program governing T-cell differentiation (**Figs. 2B-D**) and reduced TET2 expression at both the mRNA and protein levels (**Figs. 3A and 3B**). Importantly, pharmacologic inhibition of TET activity in wild-type DN thymocytes recapitulates key developmental defects observed in *Yy1^−/ΔREPO^*cells (**Figs. 5 and S4**), whereas ectopic expression of the TET2 catalytic domain partially restores T-cell development (**Fig. 6B**). These findings identify TET2 as a downstream effector of YY1 REPO domain function during early thymocyte differentiation.

Despite reduced TET2 expression, we did not detect global alterations in 5-mC or 5-hmC levels in DN thymocytes (**Fig. S1**). This observation is consistent with reports that loss of TET2 often results in locus-specific epigenetic changes without substantially altering global DNA methylation patterns, potentially due to functional compensation by TET3^11,14,65–68,81–84^. In addition, recent studies suggest that TET2 can regulate both DNA and RNA methylation, raising the possibility that multiple epigenetic mechanisms contribute to its effects on thymocyte differentiation^66,85–87^. Notably, re-expression of the TET2-CD only partially rescued the developmental defects of *Yy1^−/ΔREPO^* thymocytes (**Fig. 6B**), indicating that impaired TET2 catalytic function represents one component of a broader YY1 REPO domain-dependent regulatory program required for proper DN3 differentiation.

Our study identifies TET2 as a previously unrecognized downstream target of YY1 REPO domain-dependent regulation during early T-cell development. We found that YY1 directly binds the *Tet2* promoter (**Fig. 3C-3E**), and deletion of the YY1 REPO domain resulted in reduced *Tet2* expression at both the mRNA and protein levels in DN3 thymocytes (**Figs. 3A and 3B**). Importantly, loss of the REPO domain did not alter YY1 occupancy at the *Tet2* promoter, indicating that diminished Tet2 expression is not attributable to impaired DNA binding by YY1. Instead, *Yy1^−/ΔREPO^* DN cells exhibited reduced H3K4me3 enrichment at the *Tet2* promoter (**Fig. 3F**), consistent with a less transcriptionally permissive chromatin state. Together, these findings support a model in which the YY1 REPO domain may contribute to *Tet2* transcription by modulating the local chromatin landscape rather than through direct promoter occupancy.

This mechanism is notable because the YY1 REPO domain has traditionally been linked to Polycomb-dependent transcriptional repression and H3K27me3-associated chromatin silencing in both Drosophila and mammalian systems^38–41^. In contrast, our findings suggest that, in DN3 thymocytes, the REPO domain may facilitate maintenance of an active chromatin environment required for *Tet2* expression. This interpretation is consistent with our previous publication that YY1 deficiency in fetal liver hematopoietic stem and progenitor cells leads to globally reduced chromatin accessibility^43^. Similarly, our unpublished studies indicate that deletion of either YY1 or its REPO domain predominantly results in reduced chromatin accessibility accompanied by decreased gene expression in hematopoietic stem and progenitor cells. Collectively, these observations support an emerging model in which YY1 and its REPO domain function in a context-dependent manner to regulate chromatin accessibility and structure, acting not only as mediators of Polycomb-associated repression but also as facilitators of transcriptionally active chromatin states at selected genomic loci.

Although the molecular basis of this activity remains to be determined, our data raise the possibility that the YY1 REPO domain promotes recruitment or stabilization of chromatin regulatory complexes required to maintain H3K4me3 enrichment and transcriptional competence at the *Tet2* locus. In support of this idea, our previous studies in hematopoietic stem and progenitor cells demonstrated physical interactions between YY1 and the cohesin component SMC3, as well as extensive genomic co-localization of YY1 with SMC3 and CTCF^36^. These findings raise the possibility that YY1 may facilitate recruitment or stabilization of cohesin-and/or CTCF-associated chromatin architecture at the *Tet2* locus, thereby promoting a chromatin environment permissive for *Tet2* transcription. Future studies examining the occupancy and functional contributions of cohesin and CTCF at the *Tet2* locus in developing thymocytes will be important to test this model.

## Acknowledgments

We extend our gratitude to the University of Wisconsin Carbone Comprehensive Cancer Center (UWCCC) for providing access to the Shared Services, including the Flow Cytometry Core and Cancer Informatics Shared Resources, which are supported by UWCCC grant P30 CA014520. This work was also supported by grants TL1TR002375 and UL1TR002373 awarded to the University of Wisconsin-Madison Institute for Clinical and Translational Research (ICTR) through the National Center for Advancing Translational Sciences (NCATS). We thank Dr. Shannon Kenney for providing the pcDNA3-FLAG-TET2 plasmid and for assistance with related experimental preparation.

## Authorship Contributions

Y.W. and X.P. designed experiments. Y.W., M.N., and X.P. performed experiments. Y.W., M.N., R.L., M.B., and X.P. analyzed and interpreted the data. Y.W., R.L., M.N., and X.P. wrote the manuscript.

## Conflict of Interest Disclosures

The authors declare no competing interests.

## Notes

### Competing Interest Statement

The authors have declared no competing interest.

### Summary of Updates

Figure 2 is revised. Updated Figure 3E and 3F. Original Figure 5 was separated into Figure 5 and Figure 6.

