## Supplemental file for "Yin Yang 1-dependent TET2 regulation is essential for early T cell development"

### Supplementary Methods

#### Western blots

Antibodies detecting YY1 (ab109237; Abcam; 1:5,000), β-actin (A5441, Sigma-Aldrich, 1:10,000), TET2 (D6C7K; Cell Signaling Technology; 1:1,000) and Vinculin (E1E9V; Cell Signaling Technology; 1:1,000) proteins were used according to the manufacturer's recommendation. The intensity of specific protein bands was quantified using Adobe Photoshop software.

#### Quantitative real-time PCR

Total RNA was isolated using the RNeasy Mini Kit (Qiagen) according to the manufacturer's protocol. Complementary DNA (cDNA) was synthesized with the SuperScript III First-Strand Synthesis System Kit (Invitrogen™) with oligo (dT) primers and was subjected to the reverse transcriptase-polymerase chain reaction (RT-PCR) assay. *Yy1, Tet1, Tet2, TET2-CD,* and *Tet3* transcripts were detected by Roche LightCycler 96 with the cycle setting at 95°C (10 min), 95°C (10 s), 60°C (10 s) and 72°C (20 s) for a total of 40 cycles. *Tbp* was used as an internal control for normalization, and quantification was determined by the delta-delta cycle threshold (ΔΔCt) method.

#### Flow cytometry analysis

Directly conjugated or biotin-conjugated antibodies specific for the following surface antigens were purchased from eBioscience: CD4 (RM4-5), CD8 (53-6.7), B220 (RA3-6B2), Ter 119 (TER-119), Gr1 (RB6-8C5), CD19 (eBio1D3), IL7Ra (A7R34), Mac1 (M1/70), CD25 (PC61.5) and

CD44 (IM7), TCRβ (H57-597). All antibodies were diluted at 1:200 ratio, unless otherwise noted. Ghostdye™ Violet510 (Tonbo Bioscience) or DAPI (Thermo Fisher Scientific) were used to exclude non-viable cells. Acquisitions were performed on an LSR II Fortessa (BD Biosciences) or Cytek Aurora (Cytek Biosciences). Data were analyzed using FlowJo v10.0.7 software.

**Apoptosis assays**

Thymocytes or DN T cells from OP9-DL1 co-culture system were harvested and stained with antibodies against lineage markers (CD4, CD8, B220, CD11b, NK1.1, CD19, Ter119, and Gr-1), CD25, and CD44 to identify DN subsets. Then, cells were resuspended in the binding buffer solution with PE anti-annexin V (559763; BD Biosciences; 1:40). Next, cells were resuspended in DAPI solution (Thermo Fisher Scientific, 1:1000). Samples were analyzed using the BD LSRII Fortessa and results were analyzed using FlowJo v10.0.7.

**Ki67 proliferation assay**

Freshly isolated thymocytes were fixed with 4% paraformaldehyde, permeabilized with 0.1% saponin in PBS, and stained with PE-Ki67 (BD Biosciences, 1:40) and DAPI solution (Thermo Fisher Scientific, 1:250). Cell cycles were analyzed within DN1 ($Lin^{-}CD44^{+}CD25^{-}$), DN2 ($Lin^{-}CD44^{+}CD25^{+}$), DN3 ($Lin^{-}CD44^{-}CD25^{+}$), and DN4 ($Lin^{-}CD44^{-}CD25^{-}$) populations as previously described (40).

**RNA-seq**

DN3 thymocytes ($Lin^{-}CD25^{+}CD44^{-}$) were sorted from total thymocytes using FACS sorter. Total RNA was isolated from DN3 T cells using the RNeasy Mini Kit (Qiagen) according to the manufacturer's protocol. Sequencing libraries were prepared by using Takara SMART-Seq V4 Ultra-Low Input according to the manufacturer’s specifications and were sequenced by an Illumina NovaSeq 6000 at the University of Wisconsin Biotechnology Center Gene Expression Center. RNA-seq reads were aligned by STAR (version 2.5.2b) to the mouse genome (version mm10) with GENCODE basic gene annotations (version M25). Gene expression levels were quantified by featureCounts (Galaxy Version 2.1.1), and differential expressions were analyzed by edgeR (version 3.36.0). Differentially expressed genes were defined as genes with an absolute log2 fold change $\geq 1$, an adjusted P value $< 0.05$, and TPM $\geq 1$ in all biological replicates of at least one cohort in the comparison. GSEA was performed by fgsea (version 1.20.0) with gene sets from the Molecular Signatures Database.

**ELISA**

Genomic DNA was extracted from sorted DN3 (Lin$^-$CD44$^-$CD25$^+$) thymocytes using DNeasy Blood & Tissue Kit (Qiagen). 30 ng of gDNA was used per well for ELISA reactions. EpigenTekMethylFlash™ Global DNA Methylation (5-mC) ELISA Easy Kit (P-1030) and Global DNA Hydroxymethylation (5-hmC) ELISA Easy Kit (P-1032) were used to measure the global DNA methylation and demethylation percentage, respectively. The absorbance was measured at 450 nm by a microplate reader.

**5-hmC dot blot**

100 ng genomic DNA isolated from Lin$^-$ thymocytes (Qiagen DNeasy Blood & Tissue Kit) was dotted onto a Nylon membrane. The spotted membrane was subsequently air-dried for 10 min and UV-crosslinked at 120000 μJ/cm2. The membrane was then blocked in 5% non-fat dry milk in

Tris-buffered saline with 0.1% Tween-20 (TBS-T) for 1 h and incubated overnight at 4°C with 5-Hydroxymethylcytosine (5-hmC) antibody (pAb) (Active Motif 39092, 1:1000 diluted in 1% BSA in TBS-T). Membranes were then washed 3 times with TBS-T and blocked with a secondary antibody. The signal was detected by enhanced chemiluminescence with the Clarity Western ECL substrate.

### Supplementary Figures

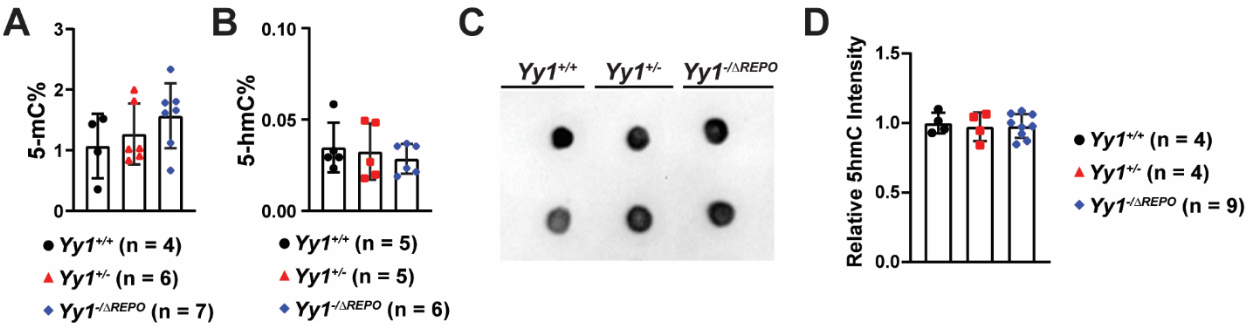


**Figure S1. Deletion of the YY1 REPO domain does not impact global DNA 5-mC and 5-hmC in DN3 T cells. (A-B)** ELISA quantification of global 5-mC% of total DNA **(A)** and 5-hmC% of total DNA **(B)** in DN3 T cells. **(C)** Representative dot blot of 5-hmC level in genomic DNA from Lin$^-$ thymocytes. **(D)** Quantification of relative 5-hmC intensity of all dot blot samples. N represents the number of mice; data are presented as means ± SD; no statistical significance was detected using one-way ANOVA.

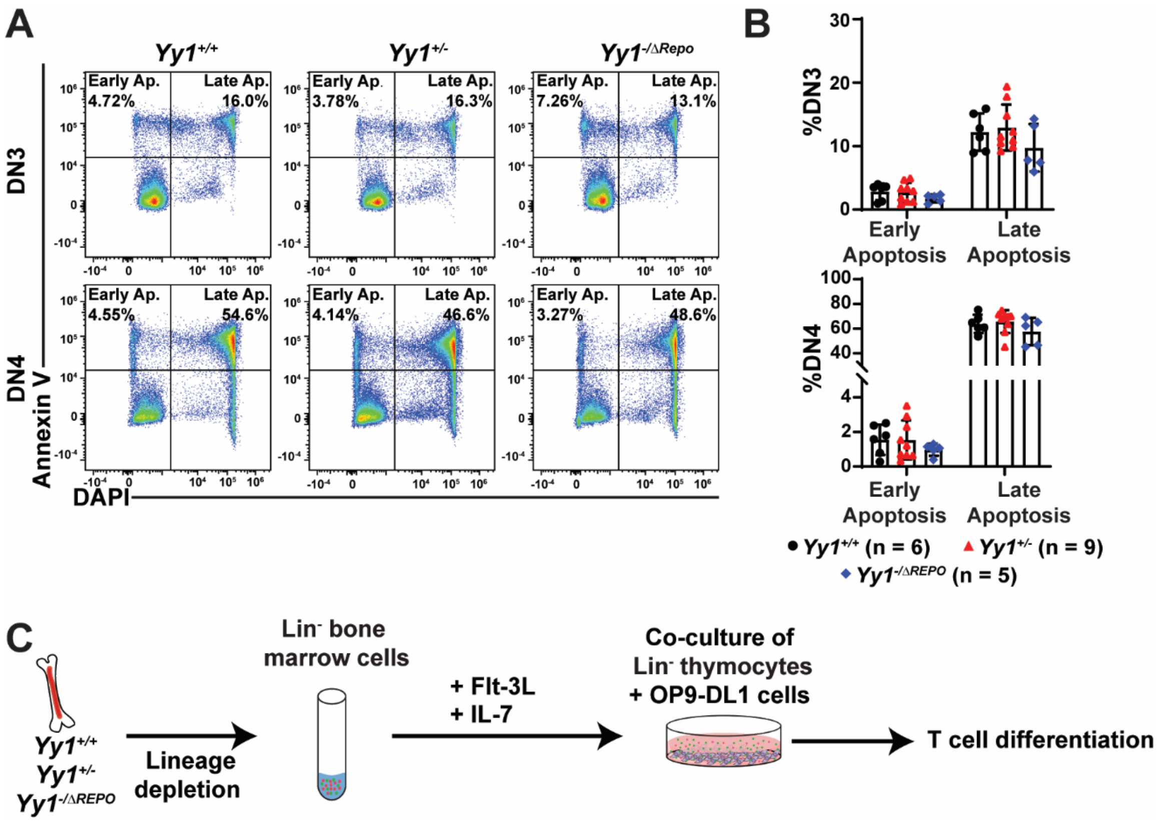


**Figure S2. In vivo survival assay did not reveal differences in apoptosis in *Yy1*$^{+/+}$, *Yy1*$^{+/-}$, and *Yy1*$^{-/\Delta REPO}$ DN3 and DN4 cells. (A)** Representative flow plots of apoptosis assay. **(B)** Quantification of early apoptotic (Annexin V$^{+}$, DAPI$^{-}$) and late apoptotic (Annexin V$^{+}$, DAPI$^{+}$) cells in DN3 and DN4 T cells. **(C)** Schematic diagram of OP9-DL1 co-culture system for in vitro apoptosis assay. Lin-BM cells from *Yy1*$^{+/+}$, *Yy1*$^{+/-}$, and *Yy1*$^{-/\Delta REPO}$ were cultured with OP9-DL1 feeder cells with IL-7 and Flt-3L. N represents the number of mice; data are presented as means ± SD; no statistical significance was detected using two-way ANOVA.

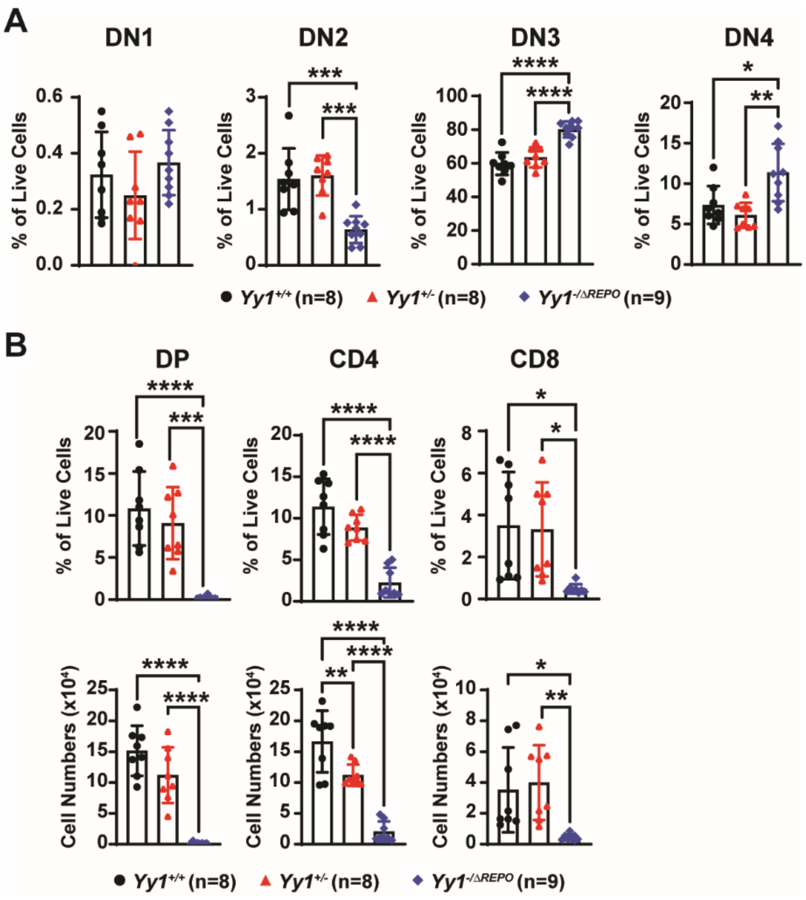


**Figure S3. The YY1 REPO deletion in OP9-DL1 co-culture system recapitulates in vivo defects.** Lin- thymocytes were plated and co-cultured with OP9-DL1 feeder cells with Flt-3L and IL-7 for 72 hours. **(A)** Percentages of DN T cells in the OP9-DL1 co-culture system. **(B)** Quantification of DP and SP T cell percentage and cell numbers. N represents the number of mice; data are presented as means ± SD; $*P < .05$, $**P < .01$, $***P < .001$, and $****P < .0001$ by one-way ANOVA.

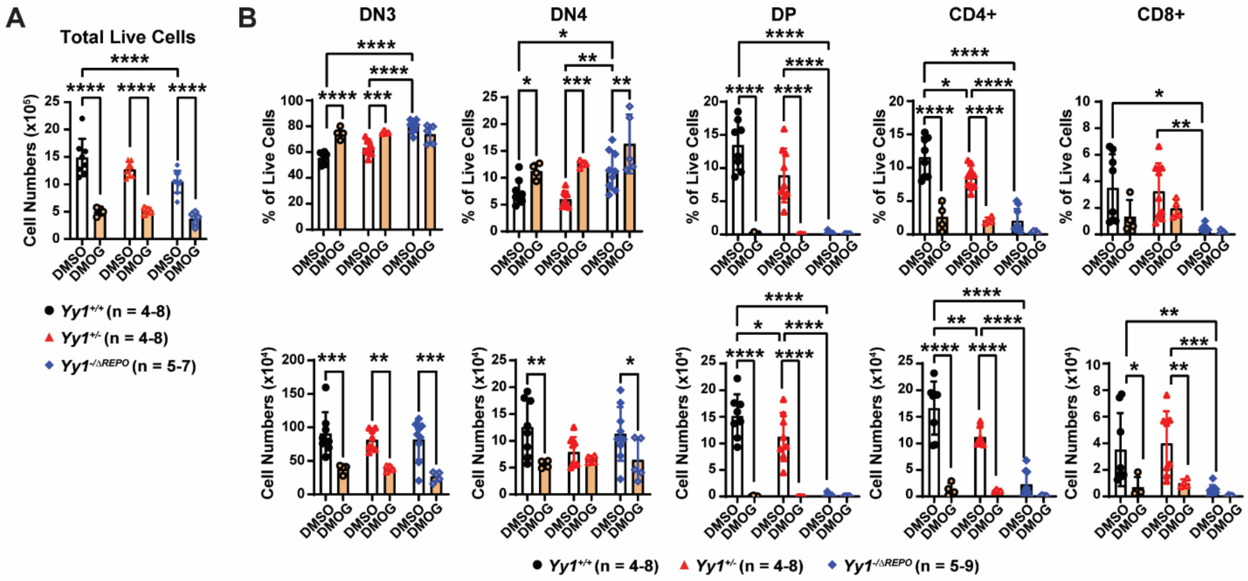


**Figure S4. Lin$^-$ thymocytes OP9-DL1 co-culture system treated with TET inhibitor DMOG.**

**(A)** Total cell counts in the co-culture system 72h after DMOG treatment or DMSO control. **(B)** Quantification of DN3, DN4, DP, CD4+, and CD8+ T cells percentage and cell numbers in the co-culture system 72h after DMOG treatment or DMSO control. N represents the number of mice except otherwise indicated; data are presented as means ± SD; $*P < .05$, $**P < .01$, $***P < .001$, and $****P < .0001$ by two-way ANOVA.

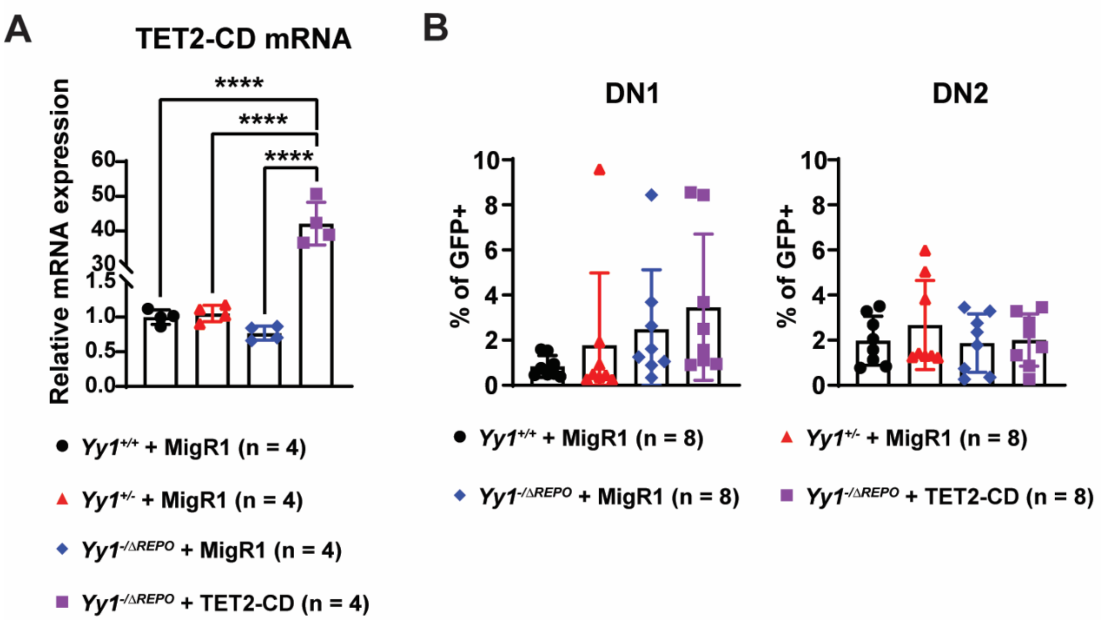


**Figure S5. TET2-CD expression in the rescue experiment. (A)** *Tet2-CD* mRNA expression in OP9-DL1 co-cultured cells transduced with MigR1 or MigR1-TET2-CD. **(B)** Percentages of DN1 and DN2 T cells at 48h after transduction with MigR1 or MigR1-TET2-CD. N represents the number of mice; data are presented as means ± SD; $*P < .05$, $**P < .01$, and $****P < .0001$ by one-way ANOVA.

**Table S1. Primers used for mRNA detection.**

| Target | Forward | Reverse |
|---|---|---|
| *Yy1* | TCAGACCCTAAGCAACTGGCAGAA | TTGAGCTCTCAACGAACGCTTTGC |
| *Tet1* | ATTTCCGCATCTGGGAACCTG | GGAAGTTGATCTTTGGGGCAAT |
| *Tet2* | TGCTTTCCCAACACGGAACTA | GCACCATTAGGCATTAGCACAAT |
| *Tet3* | GCTTCTTAAGGAGACAGGCTCAGA | TGAGGTGCTTAGCTGCCTTG |
| *Tet2-CD* | TGAGATCCTGGTGGGGTGAT | ATCCTTGCATTGGAGGGGTG |
| *Tbp* | CTACCGTGAATCTTGGCTGTAAAC | AATCAACGCAGTTGTCCGTGGC |

**Table S2. Primers used for CUT&Tag qPCR analysis.**

| Target | Forward | Reverse |
|---|---|---|
| *Rpl30* | GCAGAGCTCACTCACCAACA | GCTCCTTCCTTTCTCGCTCC |
| *Tet2* | CAAATCCTACAGGGCAGCCA | ATATTGATGCGGAGGCGAGG |
| *Gene Desert* | TCCTCCCCATCTGTGTCATC | GGATCCATCACCATCAATAACC |
